# Pro-inflammatory chemokines CCL5, CXCL12, and CX3CL1 bind to and activate platelet integrin αIIbβ3 in an allosteric manner

**DOI:** 10.1101/2022.09.01.506262

**Authors:** Yoko K Takada, Masaaki Fujita, Yoshikazu Takada

## Abstract

Previous studies showed that pro-inflammatory chemokines CX3CL1 and CXCL12 bound to the allosteric binding site (site 2) of integrins and allosterically activated integrins, in addition to the classical ligand-binding site (site 1). We showed that CCL5 also bound to site 2, in addition to site 1, and activated soluble integrin αvβ3. Platelet integrin αIIbβ3, a fibrinogen receptor, is critical for hemostasis and thrombus formation and activation of αIIbβ3 is a key event for thrombus formation. Activation of αIIbβ3 is known to be mediated exclusively by inside-out signaling. We studied if αIIbβ3 can be allosterically activated. We discovered that CCL5, CXCL12, and CX3CL1 are new ligands for αIIbβ3. Notably they enhanced the binding of monovalent ligand to soluble αIIbβ3 in 1 mM Ca^2+^ by binding to site 2. They activated cell-surface αIIbβ3 on CHO cells quickly (half maximal response <1 min) and at low concentrations (1-10 ng/ml) compared to soluble αIIbβ3, probably because chemokines bind to cell surface proteoglycans. Notably, activation of αIIbβ3 by the chemokines was several times more potent than 1 mM Mn^2+^. Since CCL5 and CXCL12 are stored in platelet granules and rapidly transported to the surface upon platelet activation, we hypothesized that they are released from the granules and allosterically activate αIIbβ3 by binding to site 2.

Transmembrane CX3CL1 on activated endothelial cells likely mediates platelet-endothelial interaction by binding to and activating αIIbβ3. Also, over-produced chemokines during inflammation may trigger αIIbβ3 activation, which is a possible missing link between inflammation and thrombosis.

## Introduction

Integrins are superfamily of cell-surface heterodimers that recognize extracellular matrix (ECM), cell-surface ligands (e.g., VCAM-1 and ICAM-1), and soluble ligands (e.g., growth factors such as FGF and IGF) (1–3). Integrins play important roles in normal biological process (e.g., wound healing and hemostasis) and in the pathogenesis of diseases (4). By virtual screening of protein data bank using docking simulation, we identified the chemokine domain of CX3CL1 (5) as a new ligand for integrins αvβ3, α4β1, and α5β1, and that CX3CL1 and integrins simultaneously bound to CX3CR1 (5). The CX3CL1 mutant defective in integrin binding was defective in signaling and acted as an antagonist although it still bound to CX3CR1 (5), indicating that the direct integrin binding to CX3CL1 and subsequent integrin-CX3CL1-CX3CR1 ternary complex formation is required for its signaling functions. CX3CL1 is expressed on the cell surface of IL-1β– and TNFα–activated endothelium as a membrane-bound form (6). Soluble CX3CL1 is released by metalloproteinases ADAM10 and ADAM17 (7–9). Interaction between membrane-bound CX3CL1 and CX3CR1 promotes leukocyte adhesion to the endothelium (10–12).

We unexpectedly discovered that CX3CL1 activated soluble integrin αvβ3 in 1 mM Ca^2+^ in cell-free conditions by binding to the second ligand-binding site (site 2), which is distinct from the classical ligand-binding site (site 1) in the integrin headpiece (13). We showed that peptides from site 2 of the integrin β subunit bound to CX3CL1 and blocked integrin activation by CX3CL1 (13). Therefore, we concluded that site 2 is the allosteric binding site and involved in allosteric activation of integrins. We showed that CXCL12 activated integrins αvβ3, α4β1, and α5β1 by binding to site 2 (14), indicating that this mechanism of integrin activation is not limited to CX3CL1. CXCL12 is a potent chemoattractant for leukocytes and is believed to regulate signaling events through CXCR4 and CXCR7 in leukocytes (15–18). CXCL12 (SDF-1) is ubiquitously expressed in many tissues and cell types. Binding of CXCL12 to CXCR4 induces trimeric G protein signaling leading to activation of the pro-inflammatory signaling (19, 20). CXCL12/CXCR4 are over-expressed in rheumatoid arthritis, systemic lupus erythematosus, multiple sclerosis, and inflammatory bowel disease (21–24). CXCL12 is stored in platelet granules and rapidly transported to the surface upon platelet activation (25).

Platelet integrin αIIbβ3 is a receptor for several proteins including fibronectin, fibrinogen, plasminogen, prothrombin, thrombospondin, vitronectin, and von Willebrand factor (VWF). Activation of αIIbβ3 is a key event that triggers platelet aggregation by inducing αIIbβ3 binding to fibrinogen leading to bridge formation between platelets (26, 27). The mechanism of integrin activation has been extensively studied in integrin αIIbβ3 as a model. Integrin activation is defined by the increase in binding to monomeric ligand and we have used monomeric fragment of fibrinogen or fibronectin.

It has been proposed that activation of integrin αIIbβ3 is induced exclusively by inside-out signaling by platelet agonists (e.g., thrombin, ADP, collagen), which bind to receptors on the cell-surface. Signals received by other receptors induce the binding of talin and kindlin to cytoplasmic end of the integrin β subunit at sites of actin polymerization (26–28). It has been, however, proposed that canonical integrin activation pathways by platelet agonists induces integrin binding to multivalent ligands (e.g., ligand-mimetic antibody, Pac-1 IgM specific for αIIbβ3, with potential 10-binding sites), but does not enhance ligand binding affinity to monovalent ligand (29, 30).

We showed that the affinity to monomeric ligands is increased by chemokines in non-αIIbβ3 integrins, but it is unclear if this occurs in platelet αIIbβ3. There is a huge gap in our knowledge here. We thus hypothesized that other chemokines that are stored in platelet granules (e.g., CCL5) are potentially involved in allosteric activation of platelet integrin αIIbβ3 and trigger thrombus formation. So, we decided to include CCL5 in the present study. It is important to study if the binding of chemokines CCL5, CX3CL1, CXCL12 to site 2 activates αIIbβ3, a model system for integrin activation.

CCL5 (Rantes) is a pro-inflammatory chemokine, recruiting leukocytes to the site of inflammation. CCL5 is chemotactic for T cells, eosinophils, and basophils, monocytes, natural-killer (NK) cells, dendritic cells, and mastocytes (31). CCL5 is mainly expressed by T-cells and monocytes (32) and it is abundantly expressed by epithelial cells, fibroblasts and thrombocytes. CCL5 is involved in numerous human diseases and disorders, including transplantations (33), anti-viral immunity (31), tumor development (34) and viral hepatitis or COVID-19 (32).

In the present study, we first describe that CCL5 bound to and activated vascular integrin αvβ3, as in CX3CL1 and CXCL12. We describe that CCL5, CXCL12, and CX3CL1 specifically bound to αIIbβ3 to site 1, indicating that they are new ligands for αIIbβ3. These chemokines activated soluble αIIbβ3 in 1 mM Ca^2+^ in cell-free conditions by binding to site 2. Also, cell-surface αIIbβ3 on CHO cells, which lack the machinery for inside-out signaling or cognate receptors for these chemokines, was more efficiently activated by these chemokines probably because they were concentrated by binding to surface proteoglycans. These findings indicate that αIIbβ3 can be activated in an allosteric manner by CXCL12 and CCL5 stored in platelet granules, and CX3CL1, which is expressed on the cell surface (e.g., activated endothelial cells). We propose a new mechanism of αIIbβ3 activation by chemokines in the platelet granules upon platelet activation. Also exogenous chemokines during inflammation (e.g., cytokines storms or autoimmune diseases) contribute to thrombus formation upon platelet activation during inflammation and is a possible missing link between inflammation and thrombosis.

## Results

### CCL5 binds to and activates integrin αvβ3

The primary goal of this study is to determine if three chemokines bind to and activate integrin αIIbβ3. Active and inactive conformations, 1L5G.pdb (open headpiece αvβ3) and 1JV2 (closed-headpiece αvβ3), are defined in αvβ3 but not in αIIbβ3. We have shown that CX3CL1 binds to the classical ligand (RGD)-binding site (site 1) in open headpiece αvβ3 and to the allosteric site (site 2) in closed headpiece αvβ3 in docking simulation (13). Also, CXCL12 bound to site 2 and activated integrins αvβ3, α4β1 and α5β1 (14). It is unclear if CCL5, a major chemokine in platelet granules, binds to and activate any integrins. Thus, we first studied if CCL5 binds to site1 and site 2 of αvβ3 before we study CCL5 binding to αIIbβ3. Our docking simulation between integrin αvβ3 and CCL5 predicted that CCL5 binds to site 1 at high affinity (docking energy −24.8 kcal/mol) (Fig. 1a) and to site 2 (docking energy −20.2 kcal/mol) (Fig. 1b). Amino acid residues predicted to be involved in site 1 and site 2 binding are shown in Tables 1 and 2.

**Fig. 1.**
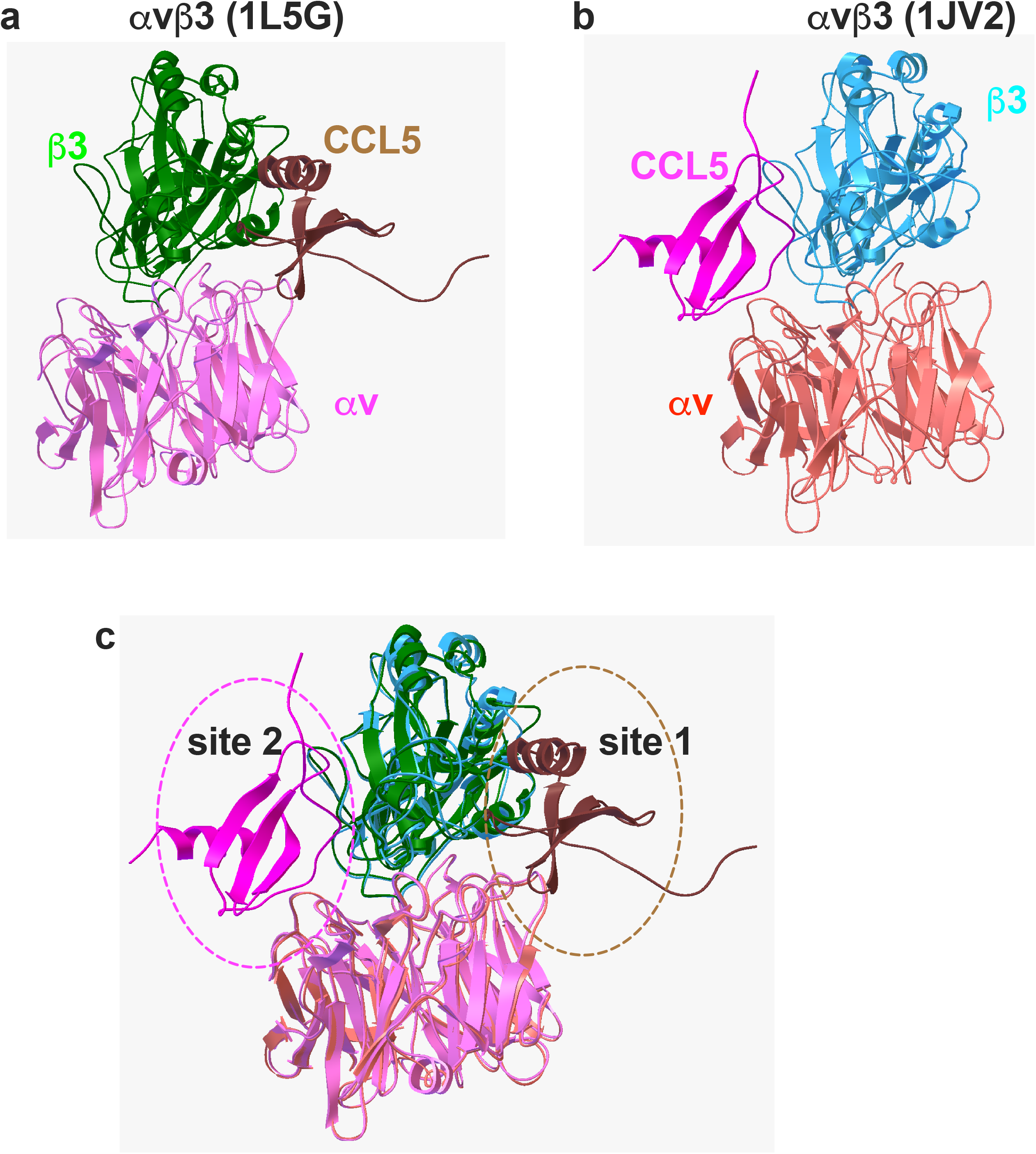
Model of the interaction between CCL5 and integrin αvβ3. Docking simulation between CCL5 (1EQT.pdb) and integrin αvβ3 (a) with open-headpiece conformation 1L5G.pdb or (b). with closed headpiece conformation, 1JV2.pdb using Autodock3, as described in methods section. The docking models were superposed (c).

**Table 1.**
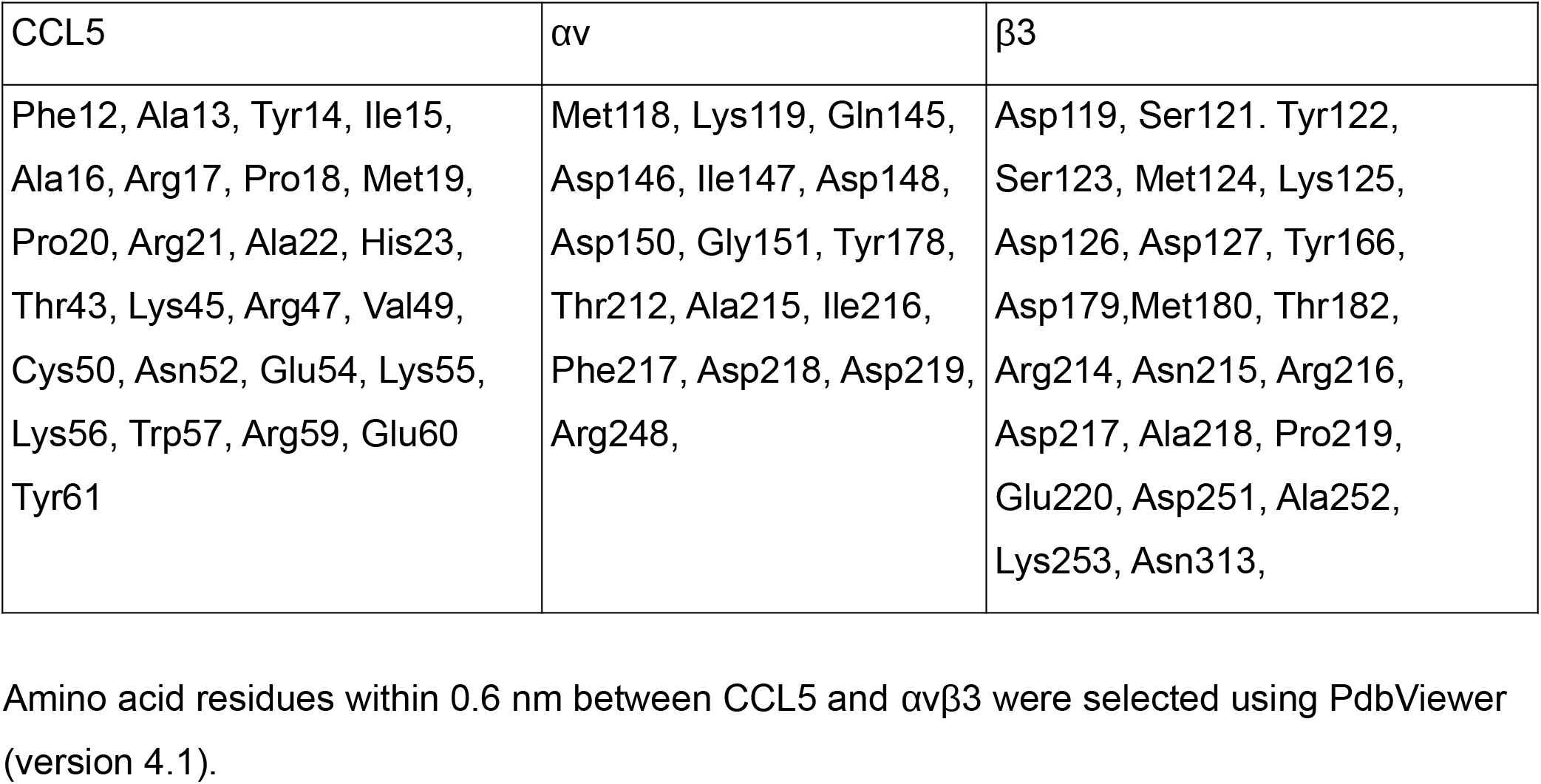
Amino acid residues involved in CCL5 (1EQT.pdb) and αvβ3 (1L5G.pdb, open headpiece) predicted by docking simulation.

**Table 2.** Amino acid residues involved in CCL5 and αvβ3 (1JV2.pdb, closed headpiece) predicted by docking simulation

| CCL5 | $\alpha\text{v}$ | $\beta 3$ |
| --- | --- | --- |
| Thr7, Thr8, Cys10, Phe12, Ala13, Tyr14, Ile15, Ala16, Arg17, Pro18, Met19, Pro20, Arg21, His23, Ser35, Asn36, Pro37, Thr43, Lys45, Arg47, Gln48, Val49, Asn52, Glu55, Lys56, Trp57, Tyr61 | Glu15, Gly16, Tyr18, Lys42, Asn44, Thr45, Thr46, Gln47, Pro48, Gly49, Ile50, Val51, Glu52, Ser90, His91 | Lys159, Pro160, Val161, Ser162, Met165, Glu171, Glu174, Asn175, Pro186, Met187, Phe188, **Val266, Gln267, Val275, Gly276, Ser277, Asp278, His280, Tyr281, Ser282, Ala283,** |
|  |  | **Thr285, Thr286** |

Amino acid residues within 0.6 nm between CCL5 and $\alpha\beta$ 3 were selected using PdbViewer (version 4.1). Amino acid residues in β3 that are in the cyclic site 2 peptide are in bold.

We studied if αvβ3 binds to CCL5 in ELISA-type binding assays. CCL5 was immobilized to wells of 96-well microtiter plate and incubated with soluble αvβ3 in 1 mM Mn^2+^. We found that soluble αvβ3 bound to immobilized CCL5 in a dose-dependent manner (Fig. 2a). The αvβ3 binding to CCL5 required Mn^2+^, and EDTA, Mg^2+^, Ca^2+^ (1 mM) did not support the binding (Fig. 2b). Inhibitory anti-αvβ3 mAb LM609 weakly but significantly suppressed the binding of soluble αvβ3 to CCL5 in 1mM Mn^2+^ (Fig. 2c), indicating that the binding of CCL5 to αvβ3 is specific. Previous studies showed that the disintegrin domain of ADAM15 (ADAM15 disintegrin) specifically bound to integrin αvβ3 (35) and later αIIbβ3 (36). We found that ADAM15 disintegrin suppressed the binding of soluble αvβ3 to the immobilized CCL5, but GST did not (Fig. 2d), indicating that CCL5 specifically bound to αvβ3. These findings indicate that CCL5 is a new ligand for αvβ3. These finding suggest that CCL5 binds to site 1 of αvβ3 in Mn^2+^-dependent manner.

**Fig. 2.**
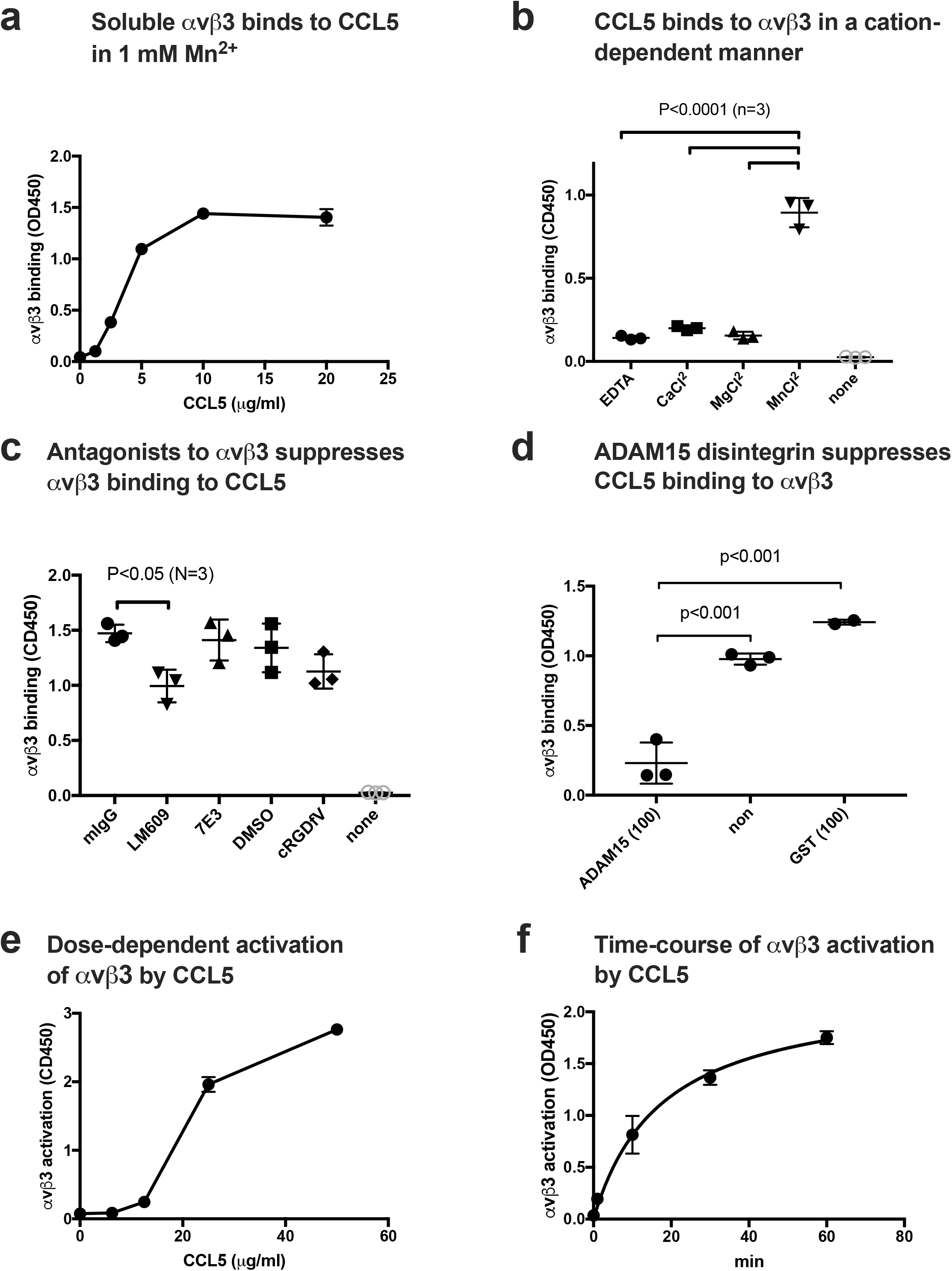
CCL5 binds to and activates integrin αvβ3. (a) <u>Binding of soluble integrin αvβ3 to CCL5 in 1 mM Mn^2+^ in cell-free conditions</u>. Wells of 96-well microtiter plate were coated with CCL5 and remaining protein-binding sites were blocked with BSA. Wells were incubated with soluble αvβ3 (1 μg/ml) in Tyrode-HEPES buffer with 1 mM Mn^2+^ (to activate αvβ3) for 1 h at room temperature. After washing the wells to remove unbound integrin, bound αvβ3 was quantified using anti-β3 mAb (AV10) and HRP-conjugated anti mouse IgG. Data are shown as means +/− SD in triplicate experiments. (b) <u>Cation dependency of αvβ3 binding to CCL5.</u> Binding of soluble integrin αvβ3 to CCL5 was measured in the presence of different cations (1 mM) in ELISA-type binding assays. Data are shown as means +/− SD in triplicate experiments. (c) <u>Effect of antagonists to αvβ3 on CCL5 binding to αvβ3.</u> The concentrations of antagonists used are 10 μg/ml for LM609 and 7E3, and 10 μM for cRGDfV. Data are shown as means +/− SD in triplicate experiments. (d) <u>ADAM15 disintegrin, a specific ligand for αvβ3, suppresses CCL5 binding to αvβ3.</u> ADAM15 disintegrin fused to GST or control GST (100 μg/ml each) were included in the binding assay as described in (a). Data are shown as means +/− SD in triplicate experiments. (e) <u>Activation of soluble αvβ3 by CCL5 in 1 mM Ca^2+^.</u> Wells of 96-well microtiter plate were coated with γC399tr, a specific αvβ3-ligand (20 μg/ml) and remaining protein-binding sites were blocked with BSA. Wells were incubated with soluble αvβ3 (1 μg/ml) in Tyrode-HEPES buffer with 1 mM Ca^2+^ for 1 h at room temperature in the presence of CCL5. After removing the unbound integrin, bound αvβ3 was quantified using anti-β3 mAb (AV10) and HRP-conjugated anti mouse IgG. Data are shown as means +/− SD in triplicate experiments. f) <u>Time-course of activation of soluble αvβ3 by CCL5.</u> Wells of 96-well microtiter plate were coated with γC399tr (a specific ligand for αvβ3) (20 μg/ml) and remaining protein-binding sites were blocked with BSA. Wells were incubated with soluble αvβ3 (1 μg/ml) and CCL5 (20 μg/ml) at room temperature in Tyrode-HEPES buffer with 1 mM Ca^2+^ for the time indicated. After washing the wells to remove unbound integrin, bound αvβ3 was quantified using anti-β3 mAb (AV10) and HRP-conjugated anti mouse IgG. Data are shown as means +/− SD in triplicate experiments.

Since docking simulation predicts that CCL5 binds to site 2 of αvβ3 (docking energy −20.2) (Fig. 1b), we studied if CCL5 activates soluble αvβ3 by binding to site 2 in ELISA-type activation assays. To study if CCL5 activate αvβ3, soluble αvβ3 was pre-incubated with CCL5 for 10 min at room temperature and then incubated with immobilized γC399tr, a specific ligand for αvβ3, in the presence of 1 mM Ca^2+^. CCL5 enhanced the binding of soluble αvβ3 to γC399tr in a dose-dependent manner (Fig. 2e), but required high concentrations (> 1 μg/ml) of CCL5 for detection. We found that CCL5 activated soluble αvβ3 in 1 mM Ca^2+^ in a time-dependent manner (Fig. 2f).

### CCL5 binds to and activates soluble αIIbβ3

Since docking simulation predicted that CCL5 binds to site 1 and site 2 of integrin αvβ3, and bound to and activated integrin αvβ3, we studied if CCL5 binds to and activates integrin αIIbβ3. We found that CCL5 bound to soluble αIIbβ3 in a dose-dependent manner (Fig. 3a). However, heat treatment (80°C 10 min) or specific inhibitors for αIIbβ3, anti-β3 mAb 7E3, or eptifibatide (0.6 μg/ml), did not affect the binding of soluble αIIbβ3 to CCL5 (data not shown). However, ADAM15 disintegrin fused to GST, a specific ligand to αIIbβ3, suppressed the binding of soluble αIIbβ3 to the immobilized CCL5, but control GST did not (Fig. 3b), indicating that CCL5 specifically bound to αIIbβ3. These findings indicate that CCL is a new ligand for αIIbβ3.

**Fig. 3.**
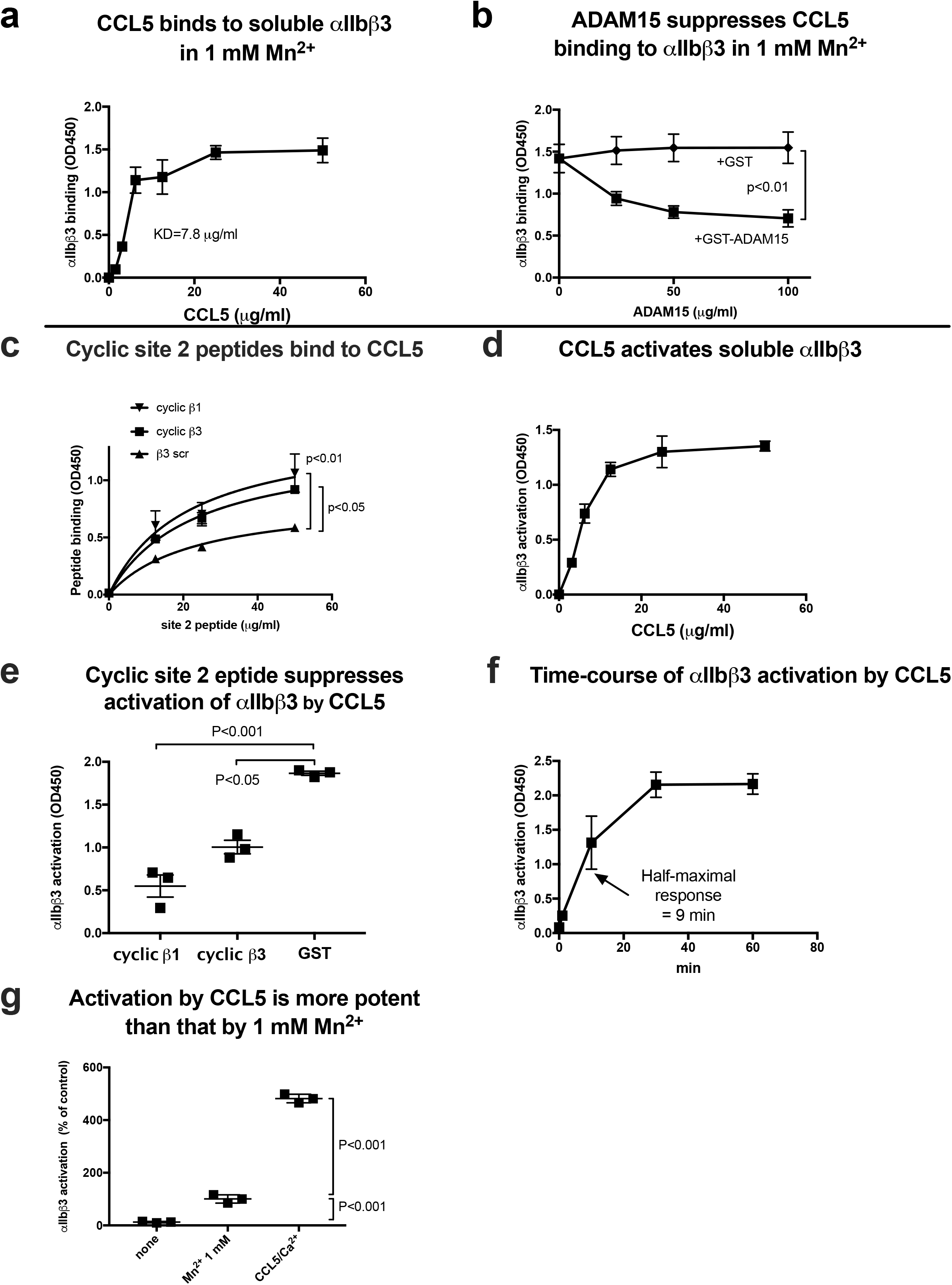
CCL5 binds to and activates soluble integrin αIIbβ3 in cell-free conditions. (a) <u>Binding of soluble integrin αIIbβ3 to CCL5 in 1 mM Mn^2+^ in cell-free conditions.</u> Wells of 96-well microtiter plate were coated with CCL5 and remaining protein-binding sites were blocked with BSA. Wells were incubated with soluble αIIbβ3 (1 μg/ml) in Tyrode-HEPES buffer with 1 mM Mn^2+^ (to activate αIIbβ3) for 1 h at room temperature. After washing the wells to remove unbound integrin, bound αIIbβ3 was quantified using anti-β3 mAb (AV10) and HRP-conjugated anti mouse IgG. Data are shown as means +/− SD in triplicate experiments. (b) <u>ADAM15 disintegrin, another ligand for αIIbβ3, suppressed CCL5 binding to αIIbβ3.</u> ADAM15 disintegrin fused to GST or control GST were included in the binding assay as described in (a). (c) <u>Binding of cyclic site 2 peptides to CCL5.</u> Wells of 96-well microtiter plate were coated with CCL5 (20 μg/ml) and remaining protein binding sites were coated with BSA. Wells were then incubated with cyclic β3 or β1 site 2 peptide fused to GST or control β3 scrambled peptide for 1 hr at room temperature and bound site 2 peptide was quantified using HRP-conjugated anti-GST antibodies. Data are shown as means +/− SD in triplicate experiments. (d) <u>Activation of soluble αIIbβ3 by CCL5 in 1 mM Ca^2+^.</u> Wells of 96-well microtiter plate were coated with γC390-411 (the αIIbβ3-ligand peptide conjugated to GST) (20 μg/ml) and the remaining protein-binding site were blocked with BSA. Wells were incubated with soluble αIIbβ3 (1 μg/ml) in Tyrode-HEPES buffer with 1 mM Ca^2+^ for 1 h at room temperature in the presence of CCL5. After removing the unbound integrin, bound αIIbβ3 was quantified using anti-β3 mAb (AV10) and HRP-conjugated anti mouse IgG. Data are shown as means +/− SD in triplicate experiments. (e) <u>Effect of site 2 peptide on αIIbβ3 activation by CCL5.</u> Activation of αIIbβ3 by CCL5 was assayed as described in (d) except that cyclic site 2 peptides or control GST (100 μg/ml each) were included as a competitor. Data are shown as means +/− SD in triplicate experiments. (f) <u>Time-course of activation of soluble αIIbβ3 by CCL5.</u> Wells of 96-well microtiter plate were coated with γC390-411 (20 μg/ml) and remaining protein-binding sites were blocked with BSA. Wells were incubated with soluble αIIbβ3 (1 μg/ml) and CCL5 (20 μg/ml) at room temperature in Tyrode-HEPES buffer with 1 mM Ca^2+^ for the time indicated. After washing the wells to remove unbound integrin, bound αIIbβ3 was quantified using anti-β3 mAb (AV10) and HRP-conjugated anti mouse IgG. Data are shown as means +/− SD in triplicate experiments. (g) <u>Comparison of αIIbβ3 activation by CCL5 and that by 1 mM Mn^2+^.</u> Soluble αIIbβ3 was activated with only 1 mM Mn^2+^ or CCL5 (50 μg/ml) in 1 mM Ca^2+^. Data are shown as means +/− SD in triplicate experiments.

Docking simulation predicts that CCL5 binds to the allosteric site (site 2) of αvβ3 at high affinity. We previously showed that peptides from site 2 bind to CX3CL1, CXCL12 and several integrin activators (e.g., sPLA2-IIA and CD40L), and blocked allosteric activation integrins αvβ3, α4β1 or α5β1 (13, 14, 37), indicating that they actually bind to site 2 and binding to site 2 is required for allosteric activation. We found that cyclic site 2 peptides from β3 and β1 bound to CCL5, indicating that CCL5 actually binds to site 2 (Fig. 3c). We thus studied if CCL5 activates soluble integrin αIIbβ3.

We previously developed ELISA-type activation assays, in which soluble integrins are incubated with ligands that are immobilized to plastic in the presence of chemokines or other activators in 1 mM Ca^2+^ (to keep integrins inactive) in cell-free conditions (13). Integrin activation is defined as the increase in binding of soluble integrins to immobilized ligand. To measure the increase in ligand binding affinity, we need to use monovalent ligand. Pac-1 IgM specific for αIIbβ3, multivalent ligand-mimetic antibody with potential 10-binding sites, has been widely used for detecting αIIbβ3 ligand binding. Since monovalent Fab of Pac-1 IgM (kindly provided by Sandy Shattil, UC San Diego, La Jolla, CA) showed very weak affinity to αIIbβ3, we generated the C-terminal residues 390-411 of fibrinogen γ-chain fused to GST (designated γC390-411) as a ligand for αIIbβ3. αIIbβ3 binds to the C-terminal 400HHLGGAKQAGDV411 sequence of fibrinogen γ-chain C-terminal domain (γC) (38). We showed that soluble recombinant αIIbβ3 (activated by 1 mM Mn^2+^) bound to γC390-411 in ELISA-type assays in a dose dependent manner (data not shown). Integrin αvβ3 does not recognize this sequence for binding to γC (39, 40). Using the whole fibrinogen γ-chain C-terminal domain (γC151-411) (39) as a ligand for soluble αIIbβ3 in the ELISA-type activation assay, we obtained results very similar to those with γC390-411 (data not shown). Since γC151-411 also binds to other integrins such as αvβ3 and αMβ2 (39, 41), we used γC390-411 as a ligand for αIIbβ3 throughout the entire project.

To study if CCL5 activate αIIbβ3, soluble αIIbβ3 was pre-incubated with CCL5 for 10 min at room temperature and then incubated with immobilized γC390-411 in the presence of 1 mM Ca^2+^. CCL5 enhanced the binding of soluble αIIbβ3 to γC390-411 in a dose-dependent manner (Fig. 3d), but required high concentrations (> 1 μg/ml) of CCL5 for detection. We found that cyclic site 2 peptides from β3 and β1 effectively suppressed CCL5-induced αIIbβ3 activation (Fig. 3e), indicating that CCL5 is required to bind to site 2 of αIIbβ3 to activate αIIbβ3. We found that it took >30 min to fully activate soluble αIIbβ3 by CCL5 (Fig. 3f). This is consistent with the previous findings that the activation of soluble integrin αvβ3 by CXCL12 was slow and required high CXCL12 concentrations (14). These findings suggest that activation of soluble αIIbβ3 by CCL5 does not require inside-out signaling, but requires high concentrations of CCL5 and takes > 60 min to complete.

It has been assumed that 1 mM Mn^2+^ fully activates integrins (42–45). We compared the levels of activation of soluble αIIbβ3 by CCL5 with that of 1 mM Mn^2+^ as a standard integrin activator. Unexpectedly, we found that CCL5 was much more potent (4.5-fold) than 1 mM Mn^2+^ (Fig. 3g).

### CXCL12 binds to and activates soluble αIIbβ3

CXCL12 is another chemokines that is stored in platelet granules and rapidly transported to the surface upon platelet activation (46). We studied if CXCL12 binds to and activates integrin αIIbβ3. We obtained results similar to that of CCL5. Briefly, CXCL12 bound to αIIbβ3 in 1 mM Mn^2+^ (Fig. 4a) and this interaction was suppressed by another αIIbβ3 ligand, ADAM15 (Fig. 4b). CXCL12 activated αIIbβ3 in a dose-dependent manner in cell-free conditions. High concentration of CXCL12 (>50 μg/ml) was required to fully activate soluble αIIbβ3 (Fig. 4c). Previous studies showed that activation of αvβ3, α4β1 or α5β1 by CXCL12 was blocked by site 2-derived peptides (14). Activation of αIIbβ3 by CXCL12 was also suppressed by cyclic site 2 peptides (Fig. 4d), indicating that CXCL12 binding to site 2 is required for αIIbβ3 activation. It took more than 60 min to fully activate soluble αIIbβ3 by CXCL12 (Fig. 4e), indicating that this activation process is relatively slow. CXCL12 was more potent (3.5-fold) in activating αIIbβ3 than that of Mn^2+^ (Fig. 4f). These findings suggest that CXCL12 also play a role in αIIbβ3 activation.

**Fig. 4.**
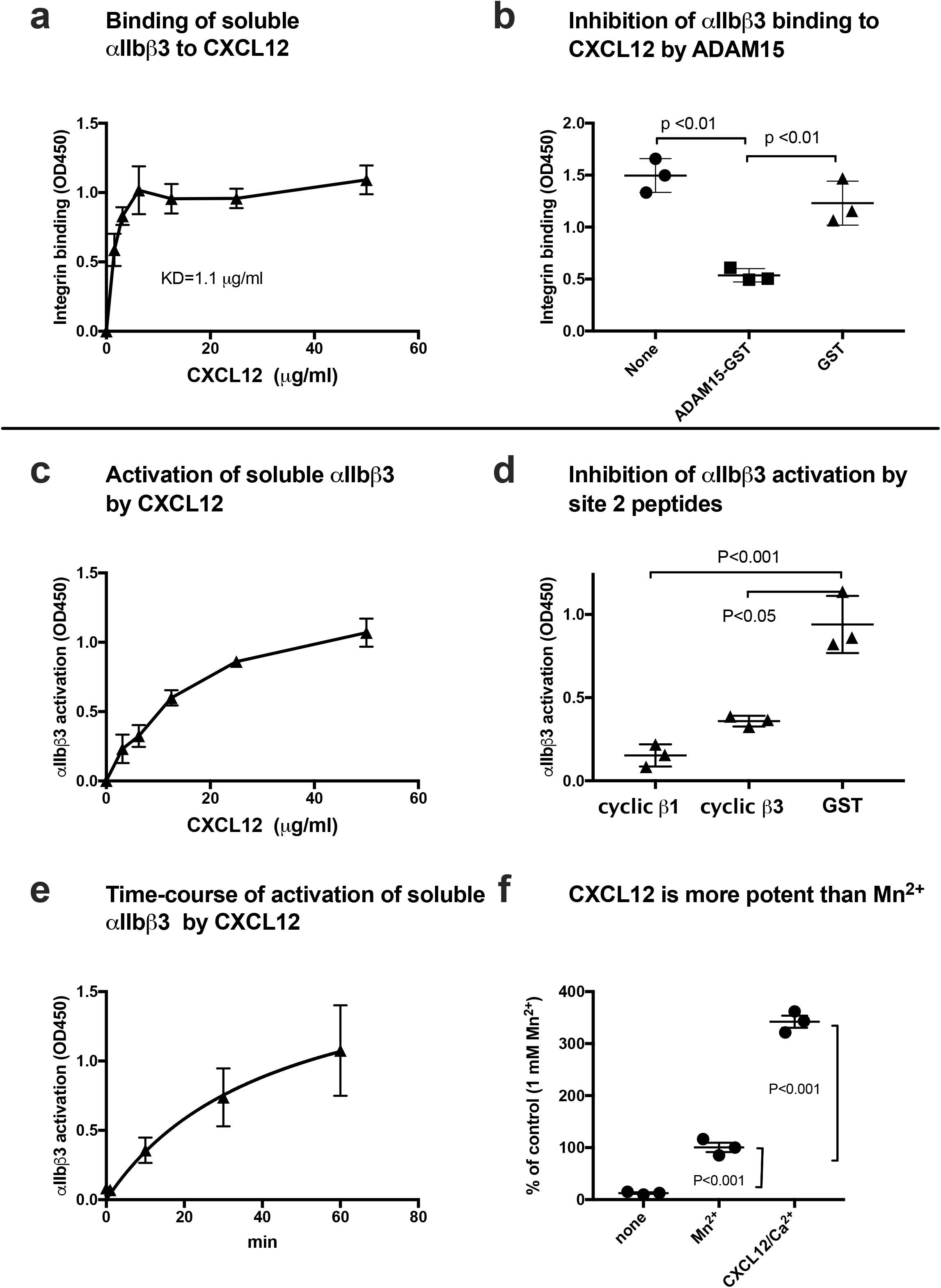
CXCL12 binds to and activates soluble integrin αIIbβ3 in cell-free conditions. (a) <u>Binding of soluble integrin αIIbβ3 to CXCL12 in 1 mM Mn^2+^ in cell-free conditions.</u> Wells of 96-well microtiter plate were coated with CXCL12 and remaining protein-binding sites were blocked with BSA. Wells were incubated with soluble αIIbβ3 (1 μg/ml) in Tyrode-HEPES buffer with 1 mM Mn^2+^ (to activate αIIbβ3) for 1 h at room temperature. After washing the wells to remove unbound integrin, bound αIIbβ3 was quantified using anti-β3 mAb (AV10) and HRP-conjugated anti mouse IgG. Data are shown as means +/− SD in triplicate experiments. (b) <u>ADAM15 disintegrin, another ligand for αIIbβ3, suppressed CXCL12 binding to αIIbβ3.</u> ADAM15 disintegrin fused to GST or control GST (100 μg/ml each) were included in the binding assay as described in (a). Data are shown as means +/− SD in triplicate experiments. (c) <u>Activation of soluble αIIbβ3 by CXCL12 in 1 mM Ca^2+^.</u> Wells of 96-well microtiter plate were coated with γC390-411 (20 μg/ml) and remaining protein-binding sites were blocked with BSA. Wells were incubated with soluble αIIbβ3 (1 μg/ml) in Tyrode-HEPES buffer with 1 mM Ca^2+^ for 1 h at room temperature in the presence of CXCL12. After removing the unbound integrin, bound αIIbβ3 was quantified using anti-β3 mAb (AV10) and HRP-conjugated anti mouse IgG. Data are shown as means +/− SD in triplicate experiments. (d) <u>Effect of site 2 peptide on αIIbβ3 activation by CXCL12.</u> Activation of αIIbβ3 by CXCL12 was assayed as described in (c) except that cyclic site 2 peptides fused to GST or control GST (100 μg/ml) were included as a competitor. Data are shown as means +/− SD in triplicate experiments. (e) <u>Time-course of activation of soluble αIIbβ3 by CXCL12.</u> Wells of 96-well microtiter plate were coated with γC390-411 (20 μg/ml) and remaining protein-binding sites were blocked with BSA. Wells were incubated with soluble αIIbβ3 (1 μg/ml) and CXCL12 (20 μg/ml) at room temperature in Tyrode-HEPES buffer with 1 mM Ca^2+^ for the time indicated. After washing the wells to remove unbound integrin, bound αIIbβ3 was quantified using anti-β3 mAb (AV10) and HRP-conjugated anti mouse IgG. Data are shown as means +/− SD in triplicate experiments. (f) <u>Comparison of αIIbβ3 activation by CXCL12 and that by 1 mM Mn^2+^.</u> Soluble αIIbβ3 was activated with only 1 mM Mn^2+^ or CXCL12 (50 μg/ml) in 1 mM Ca^2+^. Data are shown as means +/− SD in triplicate experiments.

### CX3CL1 binds to and activates soluble αIIbβ3

Transmembrane CX3CL1 is not stored in platelet granules but is expressed on the surface of activated endothelial cells (6). Previous studies showed that platelet-endothelial interaction plays a role in vascular inflammation (e.g., atherosclerosis) (25). We hypothesized that CX3CL1 on endothelial cells mediates platelet-endothelial interaction by binding to and activating αIIbβ3. We obtained results with the chemokine domain of CX3CL1 similar to that of CCL5. Briefly, we found that CX3CL1 bound to αIIbβ3 in 1 mM Mn^2+^ (Fig. 5a) and this interaction was suppressed by another αIIbβ3 ligand, ADAM15 (Fig. 5b). CX3CL1 activated soluble αIIbβ3 in ELISA-type activation assays in 1 mM Ca^2+^ and high concentrations of CXCL1 (>50 ug/ml) was required to fully activate αIIbβ3 (Fig. 5c). This activation was suppressed by cyclic site 2 peptides (Fig. 5d), indicating that CX3CL1 binding to site 2 is required for αIIbβ3 activation. It took more than 60 min to fully activate soluble αIIbβ3 by CX3CL1 (Fig. 5e), indicating that this activation process is relatively slow. CX3CL1 was several times more potent in activating αIIbβ3 than that of Mn^2+^ (Fig. 5f). These findings suggest that CX3CL1 on activated endothelial cells mediates platelet-endothelial interaction by activating and binding to platelet αIIbβ3. Also, αIIbβ3 can be activated by soluble CX3CL1 in circulation.

**Fig. 5.**
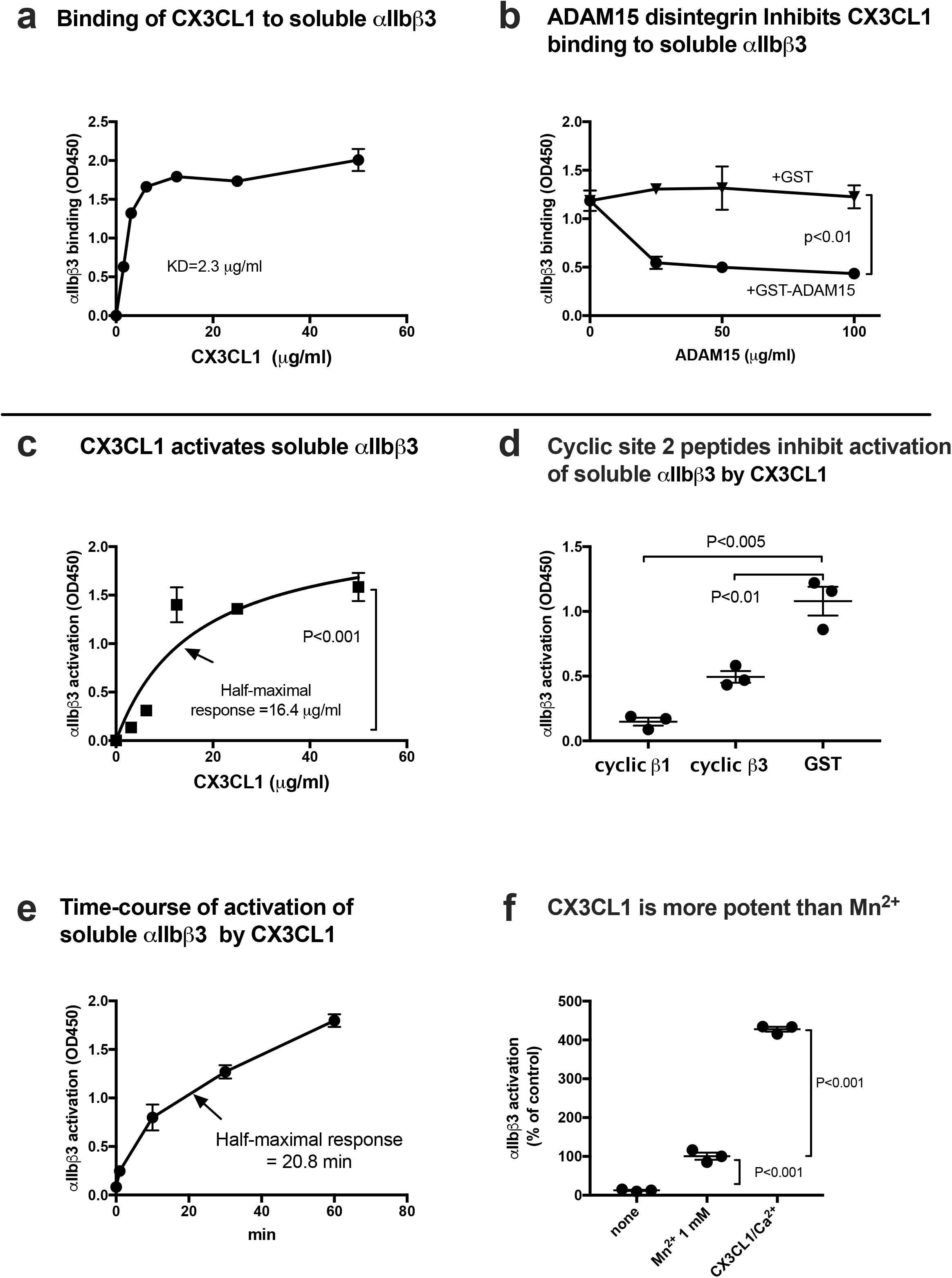
CX3CL1 binds to and activates soluble integrin αIIbβ3 in cell-free conditions. (a) <u>Binding of soluble integrin αIIbβ3 to CX3CL1 in 1 mM Mn^2+^ in cell-free conditions.</u> Wells of 96-well microtiter plate were coated with CX3CL1 and remaining protein-binding sites were blocked with BSA. Wells were incubated with soluble αIIbβ3 (1 μg/ml) in Tyrode-HEPES buffer with 1 mM Mn^2+^ (to activate αIIbβ3) for 1 h at room temperature. After washing the wells to remove unbound integrin, bound αIIbβ3 was quantified using anti-β3 mAb (AV10) and HRP-conjugated anti mouse IgG. Data are shown as means +/− SD in triplicate experiments. (b) <u>ADAM15 disintegrin suppressed CX3CL1 binding to αIIbβ3.</u> ADAM15 disintegrin fused to GST or control GST were included in the binding assay as described in (a). (c) <u>Activation of soluble αIIbβ3 by CX3CL1 in 1 mM Ca^2+^.</u> Wells of 96-well microtiter plate were coated with γC390-411 (20 μg/ml) and the remaining protein-binding site were blocked with BSA. Wells were incubated with soluble αIIbβ3 (1 μg/ml) in Tyrode-HEPES buffer with 1 mM Ca^2+^ for 1 h at room temperature in the presence of CX3CL1. After removing the unbound integrin, bound αIIbβ3 was quantified using anti-β3 mAb (AV10) and HRP-conjugated anti mouse IgG. Data are shown as means +/− SD in triplicate experiments. (d) <u>Effect of site 2 peptide on αIIbβ3 activation by CX3CL1.</u> Activation of αIIbβ3 by CXCL12 was assayed as described in (c) except that cyclic site 2 peptides (100 μg/ml) were included as a competitor. Data are shown as means +/− SD in triplicate experiments. (e) <u>Time-course of activation of soluble αIIbβ3 by CX3CL1.</u> Wells of 96-well microtiter plate were coated with γC390-411 (20 μg/ml) and the remaining protein-binding site were blocked with BSA. Wells were incubated with soluble αIIbβ3 (1 μg/ml) and CX3CL1 (20 μg/ml) at room temperature in Tyrode-HEPES buffer with 1 mM Ca^2+^ for the time indicated. After washing the wells to remove unbound integrin, bound αIIbβ3 was quantified using anti-β3 mAb (AV10) and HRP-conjugated anti mouse IgG. Data are shown as means +/− SD in triplicate experiments. (f) <u>Comparison of αIIbβ3 activation by CX3CL1 and that by 1 mM Mn^2+^.</u> Soluble αIIbβ3 was activated with only 1 mM Mn^2+^ or CX3CL1 (50 μg/ml) in 1 mM Ca^2+^. Data are shown as means +/− SD in triplicate experiments.

**Fig. 6.**
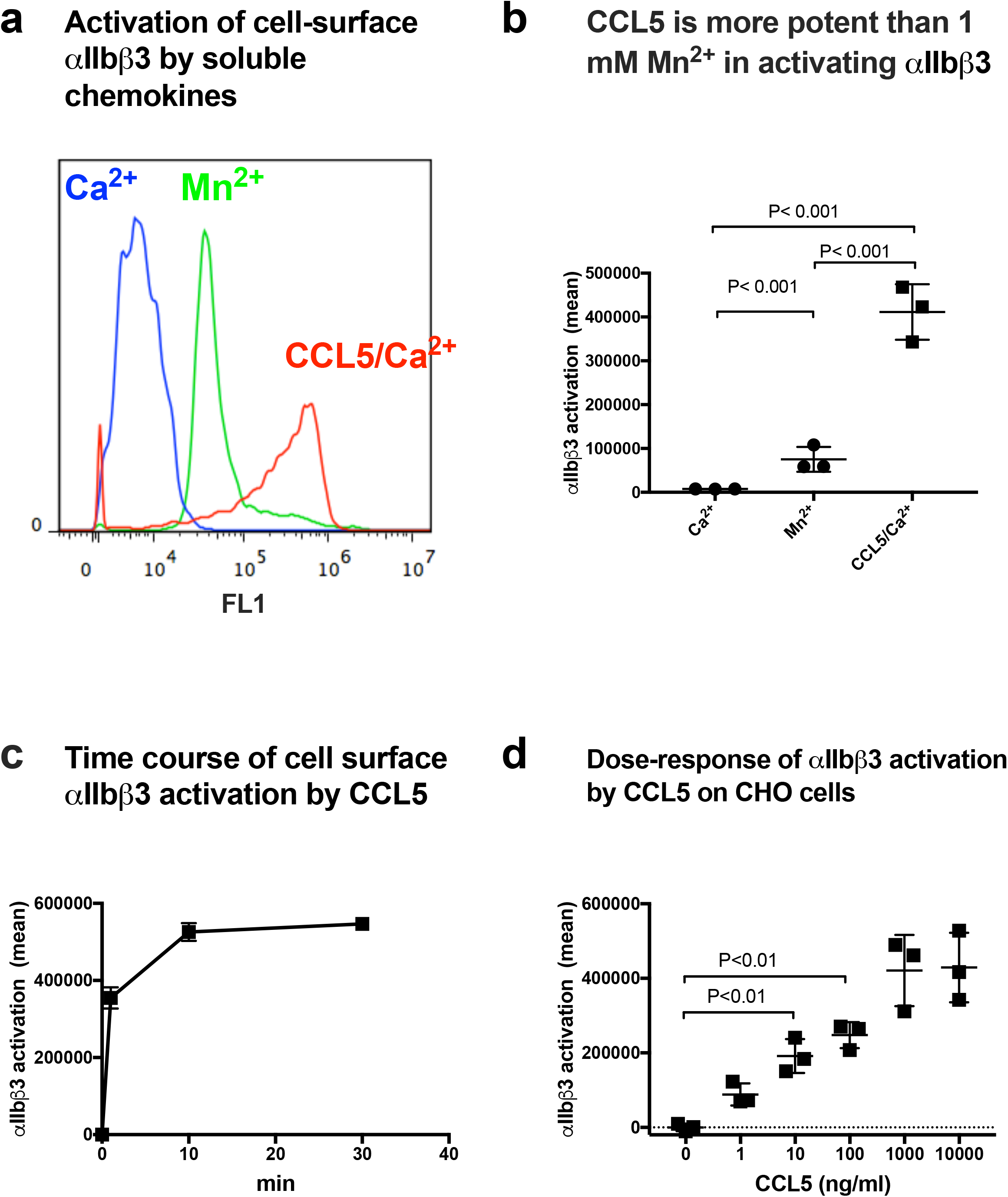
CCL5 activates cell-surface αIIbβ3 in 1 mM Ca^2+^. (a) <u>Activation of cell-surface αIIbβ3 on CHO cells.</u> αIIbβ3-CHO cells were incubated with chemokines (50 μg/ml) for 30 min on ice and then incubated with FITC-labeled γC390-411 for 30 min at room temperature. The cells were washed with PBS/0.02% BSA and analyzed by flow cytometer. Data are shown as mean fluorescence intensity +/− SD in triplicate experiments. (b) <u>Comparison of activation of cell-surface αIIbβ3 by 1 mM Mn^2+^ and chemokines in 1 mM Ca^2+^.</u> αIIbβ3-CHO cells were incubated with chemokines as described in (a). Data are shown as mean fluorescence intensity +/− SD in triplicate experiments. Ca^2+^ only and Mn^2+^ only are significantly different in unpaired t test (n=3), P< 0.001. (c) <u>Time-course of activation of cell-surface αIIbβ3 on αIIbβ3-CHO cells.</u> αIIbβ3-CHO cells were incubated with or CCL5 (50 μg/ml) and FITC-labeled γC390-411 and incubated for the time indicted at room temperature. The cells were washed with PBS/0.02% BSA and analyzed by flow cytometer. Data are shown as mean fluorescence intensity +/− SD in triplicate experiments. (d) <u>Dose-response of CCL5-induced activation of cell-surface αIIbβ3.</u> αIIbβ3-CHO cells were incubated with CCL5 at indicated concentrations for 30 min on ice and then incubated with FITC-labeled γC390-411 for 30 min at room temperature. The cells were washed with PBS/0.02% BSA and analyzed by flow cytometer. Data are shown as mean fluorescence intensity +/− SD in triplicate experiments.

### CCL5, CXCL12, and CX3CL1 more efficiently activate cell-surface αIIbβ3 than soluble αIIbβ3

αIIbβ3 on CHO cells is not activated by platelet agonists, since intracellular machineries for inside-out activation are missing (47). Previous studies showed that integrin activation by CX3CL1 or CXCL12 in CHO cells is independent of their receptors (CX3CR1 and CXCR4, respectively) (13, 14). CHO cells do not express CCR3 (48). Also, response to CCL5 in CHO cells required transfection of CCR1 or CCR5 (49, 50). Therefore, it is highly likely that activation of integrin αIIbβ3 by CCL5, CXCL12, and CX3CL1 in CHO cells occurs independent of their cognate receptors. We studied if CCL5, CXCL12, and CX3CL1 activate cell-surface αIIbβ3 on CHO cells using FITC-labeled γC390-411 in 1 mM Ca^2+^ in flow cytometry. We found that CCL5, CXCL12, or CX3CL1 markedly enhanced the binding of γC390-411 to cell-surface αIIbβ3 (Figs. 7a, 8a, and 9a).

**Fig. 7.**
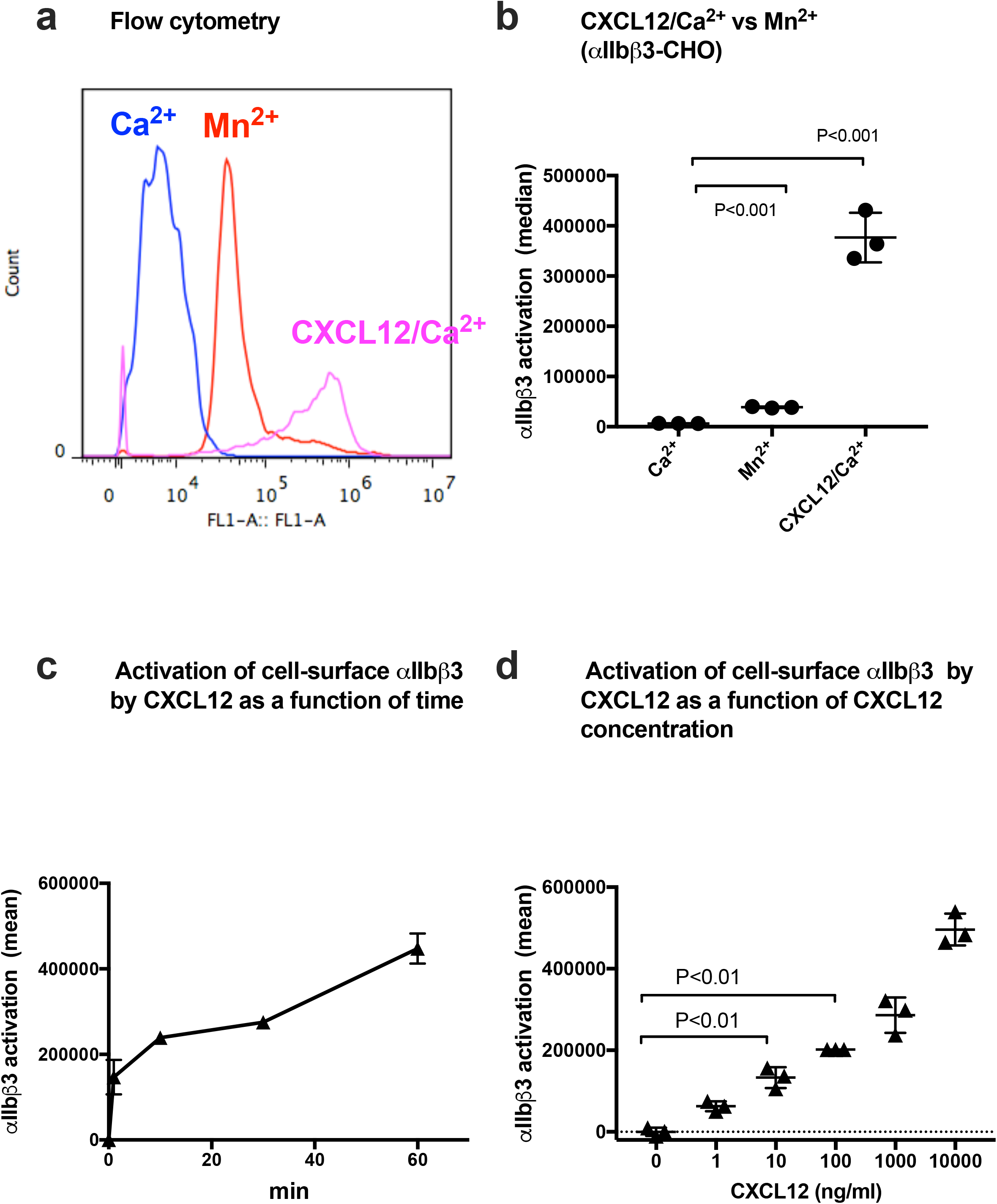
CXCL12 activates cell-surface αIIbβ3 in 1 mM Ca^2+^. (a) <u>Activation of cell-surface αIIbβ3 on CHO cells.</u> αIIbβ3-CHO cells were incubated with chemokines (50 μg/ml) for 30 min on ice and then incubated with FITC-labeled γC390-411 for 30 min at room temperature. The cells were washed with PBS/0.02% BSA and analyzed by flow cytometer. Data are shown as mean fluorescence intensity +/− SD in triplicate experiments. (b) <u>Comparison of activation of cell-surface αIIbβ3 by 1 mM Mn^2+^ and chemokines in 1 mM Ca^2+^.</u> αIIbβ3-CHO cells were incubated with chemokines as described in (a). Data are shown as mean fluorescence intensity +/− SD in triplicate experiments. Ca^2+^ only and Mn^2+^ only are significantly different in unpaired t test (n=3), P< 0.001. (c) <u>Time-course of activation of cell-surface αIIbβ3 on αIIbβ3-CHO cells.</u> αIIbβ3-CHO cells were incubated with CXCL12 (50 μg/ml) and FITC-labeled γC390-411 and incubated for the time indicted at room temperature. The cells were washed with PBS/0.02% BSA and analyzed by flow cytometer. Data are shown as mean fluorescence intensity +/− SD in triplicate experiments. (d) <u>Dose-response of CXCL12-induced activation of cell-surface αIIbβ3.</u> αIIbβ3-CHO cells were incubated with CXCL12 at indicated concentrations for 30 min on ice and then incubated with FITC-labeled γC390-411 for 30 min at room temperature. The cells were washed with PBS/0.02% BSA and analyzed by flow cytometer. Data are shown as mean fluorescence intensity +/− SD in triplicate experiments.

**Fig. 8.**
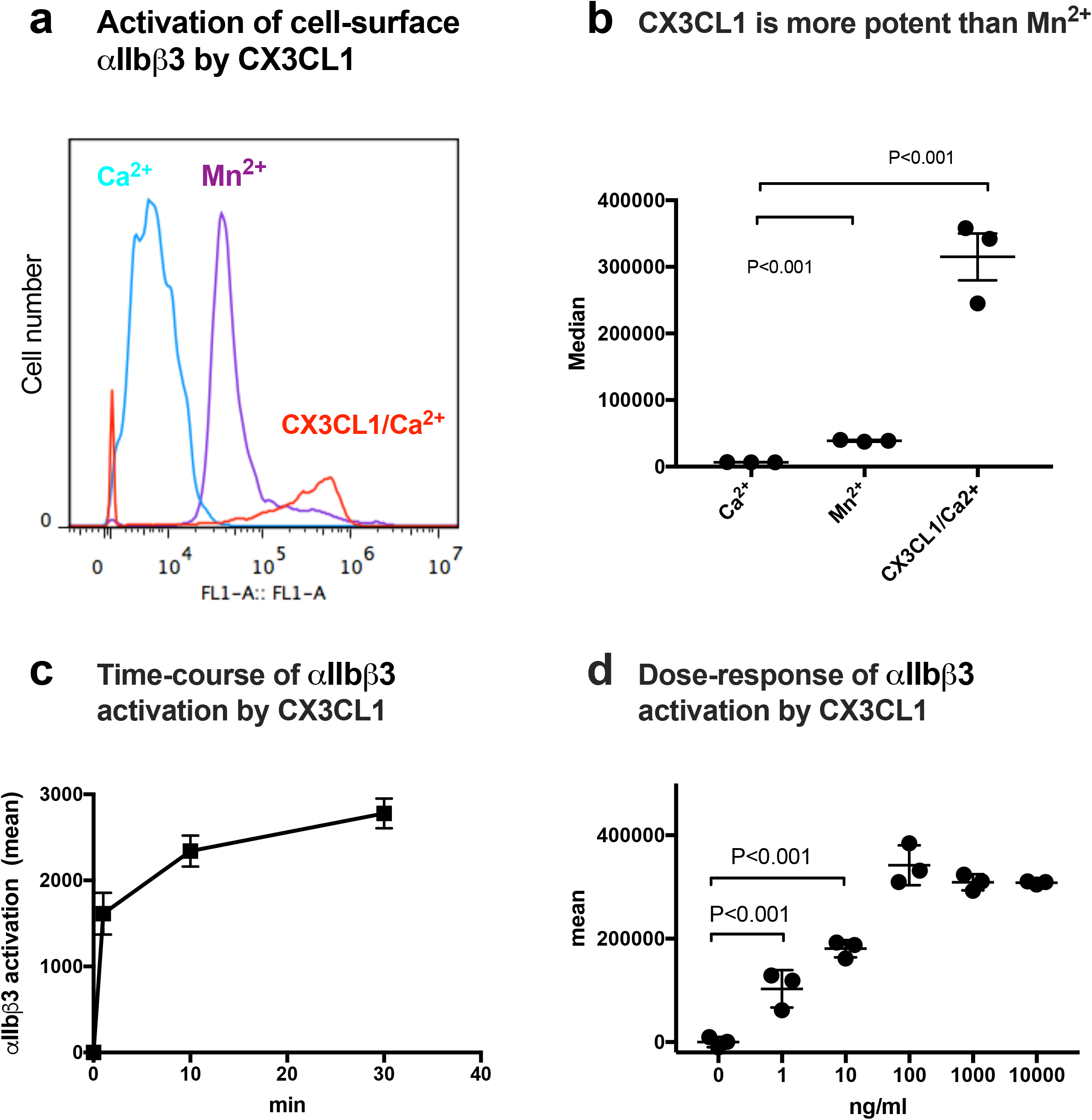
CX3CL1 activates cell-surface αIIbβ3 in 1 mM Ca^2+^. (a) <u>Activation of cell-surface αIIbβ3 on CHO cells</u>. αIIbβ3-CHO cells were incubated with chemokines (50 μg/ml) for 30 min on ice and then incubated with FITC-labeled γC390-411 for 30 min at room temperature. The cells were washed with PBS/0.02% BSA and analyzed by flow cytometer. Data are shown as mean fluorescence intensity +/− SD in triplicate experiments. (b) <u>Comparison of activation of cell-surface αIIbβ3 by 1 mM Mn^2+^ and chemokines in 1 mM Ca^2+^.</u> αIIbβ3-CHO cells were incubated with chemokines as described in (a). Data are shown as mean fluorescence intensity +/− SD in triplicate experiments. Ca^2+^ only and Mn^2+^ only are significantly different in unpaired t test (n=3), P< 0.001. (c) <u>Time-course of activation of cell-surface αIIbβ3 on αIIbβ3-CHO cells.</u> αIIbβ3-CHO cells were incubated with CX3CL1 (50 μg/ml) and FITC-labeled γC390-411 and incubated for the time indicted at room temperature. The cells were washed with PBS/0.02% BSA and analyzed by flow cytometer. The flow cytometer used in this experiment is not the same as those used for other experiments. Data are shown as mean fluorescence intensity +/− SD in triplicate experiments. (d) <u>Dose-response of CX3CL1-induced activation of cell-surface αIIbβ3.</u> αIIbβ3-CHO cells were incubated with CX3CL1 at indicated concentrations for 30 min on ice and then incubated with FITC-labeled γC390-411 for 30 min at room temperature. The cells were washed with PBS/0.02% BSA and analyzed by flow cytometer. Data are shown as mean fluorescence intensity +/− SD in triplicate experiments.

**Fig. 9.**
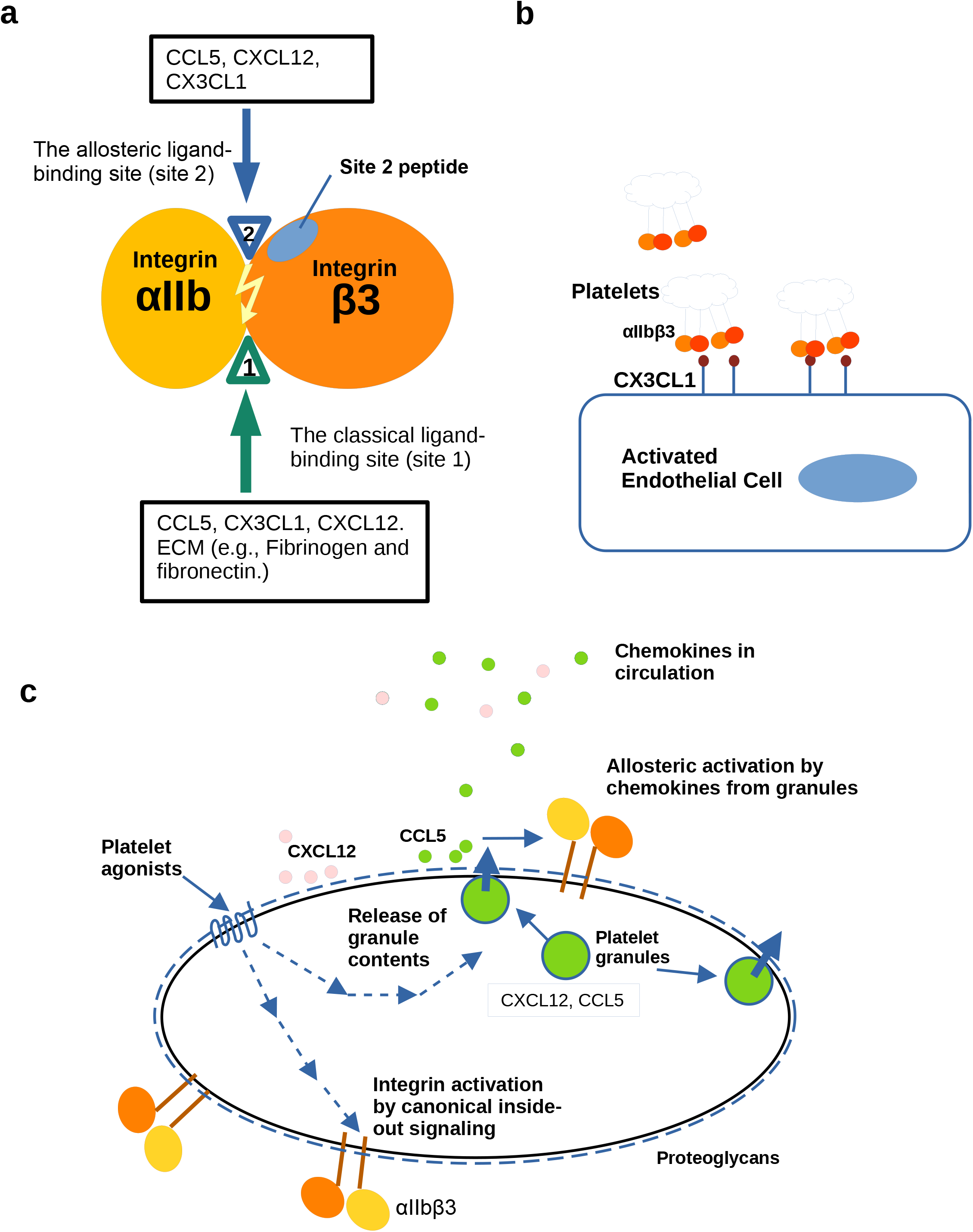
Possible biological role of allosteric activation of αIIbβ3 by CCL5, CXCL12, and CX3CL1. <u>(a) A model of the binding of CCL5, CXCL12, and CX3CL1 to the classical ligand binding site (site 1) and to the allosteric site (site 2) in integrin αIIbβ3.</u> CCL5, CXCL12, and CX3CL1 bind to site 1 and activate integrins by binding to site 2. Site 2 peptide inhibits integrin activation by chemokines. (b) <u>Potential role of CX3CL1 in platelet-endothelial cell interaction.</u> Platelet-endothelial cell interaction is involved in vascular inflammation (e.g., atherosclerosis). Transmembrane CX3CL1 is expressed on endothelial cells activated by pro-inflammatory cytokines (e.g., IL-1 β and TNFα). CX3CL1 on the cell surface can activate αIIbβ3 by binding to site 2 and then support platelet-endothelial interaction by binding to site 1. <u>(c) Activation of αIIbβ3 is induced by CXCL12 or CCL5 stored in platelet granules upon platelet activation or circulating CCL5, CXCL12 or CX3CL1 overproduced during inflammation.</u> Several chemokines are stored in platelet granules and are transported to the platelet surface upon platelet activation by platelet agonists (e.g., thrombin, collagen and ADP). Platelet integrin αIIbβ3 is inactive in resting platelets but quickly activated upon platelet activation. This activation is mediated by canonical inside-out signaling. In the present study we showed that the ligand-binding affinity of αIIbβ3 is enhanced by the binding of chemokines to site 2. Note that transfer of granular contents to the surface is mediated by inside-out signaling by platelet agonists. Levels of pro-inflammatory chemokines in circulation increase during inflammation and they may also allosterically activate αIIbβ3. This may be a missing link between inflammation and thrombosis.

We studied the time-course of activation of cell-surface αIIbβ3 by CCL5, CXCL12, and CX3CL1. CCL5, CXCL12, and CX3CL1 activated cell-surface αIIbβ3 on CHO cells much more rapidly (half-maximal response <1 min) (Figs. 7c, 8c and 9c) than in soluble αIIbβ3. Activation of soluble αIIbβ3 by CCL5, CXCL12, or CX3CL1 was slow and required more than 60 min to reach maximum (See above). Notably, the levels of activation by CCL5, CXCL12, and CX3CL1 was much higher than 1 mM Mn^2+^ (Figs. 7b, 8b, and 9b), which is consistent with the results in the ELISA-type activation assays.

We studied the activation of cell-surface αIIbβ3 as a function of CCL5, CXCL12, and CX3CL1 concentrations. We found that activation of cell-surface αIIbβ3 was detectable at 1-10 ng/ml concentrations (Figs. 7d, 8d, and 9d). We suspect that this is probably because soluble chemokines are rapidly concentrated on the cell surface by binding to cell-surface proteoglycans (14). These findings suggest that activation of platelet surface αIIbβ3 can be efficiently activated by these chemokines in an allosteric manner in the absence of canonical inside-out signaling by these chemokines, and this process is biologically relevant.

## Discussion

The present study establishes that CCL5 bound to site 1 of αvβ3, and activated αvβ3 by binding to site 2. This indicates that CCL5 is similar to CX3CL1 and CXCL12 in binding to and activating integrin αvβ3 (13, 14). Next, we showed that chemokines CCL5, CXCL12, and CX3CL1 bound to platelet integrin αIIbβ3, indicating that they are new ligands for αIIbβ3. Furthermore, activation of soluble integrin αIIbβ3 was suppressed by cyclic site 2 peptides, suggesting that the binding of these chemokines to site 2 is required. Since this activation is observed in cell-free conditions, inside-out signaling or their cognate receptors is not involved in this activation. High concentrations (>10 μg/ml) of the chemokines was required to detect activation of soluble αIIbβ3 and this activation was a slow process. These chemokines activated cell-surface αIIbβ3 on CHO cells, which lack machinery for inside-out signaling and cognate receptors for chemokines. In contrast to activation of soluble αIIbβ3, activation of cell-surface αIIbβ3 can be detected at very low concentrations of chemokines (1-10 ng/ml) and much more quickly (half maximal response < 1 min) than that of soluble αIIbβ3 probably because soluble chemokines can be quickly concentrated on the cell surface by binding to cell surface proteoglycans. These findings are the first evidence that platelet integrin αIIbβ3 can be efficiently activated by chemokines independent of canonical inside-out signaling. We propose that this is one of the mechanisms for enhancing ligand binding affinity of αIIbβ3, which is distinct from the canonical inside-out signaling (Fig. 9). CCL5 and CXCL12, which are stored in platelet granules, and CX3CL1 on the surface of activated endothelial cells, and those in circulation (e.g., during inflammation and cytokine storm) may contribute to αIIbβ3 activation. We propose that chemokine binding to site 2 of αIIbβ3 is a potential target for drug discovery and site 2 peptides have potential as therapeutics for thrombosis.

Since activation of integrins by binding of chemokines to site 2 (allosteric activation) enhanced the binding of monovalent ligand to integrins, we propose that site 2-mediated allosteric integrin activation enhanced ligand affinity of αIIbβ3. It is possible that inside-out signaling induces integrin binding to multivalent ligands (e.g., integrin clustering) and site 2-mediated allosteric binding enhances ligand affinity. Since transport of platelet granules and their contents to the surface is known to require inside-out signaling, allosteric activation by chemokines also requires inside-out signaling. It is prerequisite to determine the role of each pathway in the future studies.

Binding of integrins to their ligands requires the presence of divalent cations, and different cations can markedly alter integrin affinities to fibronectin (42), and Mn^2+^ produced the most striking increase in integrin ligand affinity compared with other divalent cations (Mg^2+^ or Ca^2+^) in a wide variety of integrins. Subsequently, Mn^2+^ has been widely used as a positive control for integrin activation. Mn^2+^-induced integrin activation was thought to mimic physiologic integrin activation because Mn^2+^ activates integrins in the absence of a bound ligand and induces similar epitope exposure (51). It is surprising that CX3CL1, CXCL12, and CCL5 are more effective in activating αIIbβ3 than 1 mM Mn^2+^. It is likely that αIIbβ3 is less sensitive to Mn^2+^ than other integrins. It is possible to use chemokines as standard of αIIbβ3 activation in future studies.

It is unclear if αIIbβ3 activation mediated by binding of chemokines site 2 requires global conformational changes in αIIbβ3. Previous studies showed that sPLA2-IIA allosterically activated integrins αvβ3, α4β1, and α5β1 (37). sPLA2-IIA did not change reactivity to activation-dependent anti-β1 antibodies HUTS4 and HUTS21 (52, 53). In our preliminary experiments, chemokines did not change reactivity of conformation-dependent anti-β3 integrin antibody LIBS2 (54) in 1 mM Ca^2+^ (data not shown). It is thus possible that allosteric activation of αIIbβ3 by chemokines may not induce global conformational changes as in allosteric activation of β1 integrins (37). It has recently been reported that the binding of allosteric activating anti β1 mAb TS2/16 to integrin α6β1 did not change the 3D structure of the integrin at all (55). It would be important to study if the binding of chemokines to site 2 induces changes in 3D structure of integrins in future experiments.

## Experimental Procedures

### Materials

cDNA encoding (6 His tag and Fibrinogen γ-chain C-terminal residues 390-411)[HHHHHH]NRLTIGEGQQHHLGGAKQAGDV] was conjugated with the C-terminus of GST (designated γC390-411) in pGEXT2 vector (BamHI/EcoRI site). The protein was synthesized in E. coli BL21 and purified using glutathione affinity chromatography. CHO cells that express recombinant human αIIbβ3 were described (56). The fibrinogen γ-chain C-terminal domain (γC151-411) was generated as previously described (39). CX3CL1 (5) and CXCL12 (14) were synthesized as described. cDNA encoding mature CCL5 (SPYSSDTTPCCFAYIARPLPRAHIKEYFYTSGKCSNPAVVFVTRKNRQVCANPEKKWVRE YINSLEMS) was synthesized and subcloned into the BamH1/EcoR1 site of pET28a vector. Protein synthesis was induced by IPTG in E. coli BL21. Proteins were produced as insoluble inclusion bodies and purified in denaturing conditions and refolded as described (5). The disintegrin domain of ADAM15 fused to GST (ADMA15 disintegrin) and parent GST were synthesized as previously described (35).

Cyclic β3 site 2 peptide fused to GST-The 29-mer cyclic β3 site 2 peptide C260-RLAGIV[QPNDGSHVGSDNHYSASTTM]C288 was synthesized by inserting oligonucleotides encoding this sequence into the BamHI-EcoRI site of pGEX-2T vector. The positions of Cys residues for disulfide linkage were selected by using Disulfide by Design-2 (DbD2) software (http://cptweb.cpt.wayne.edu/DbD2/) (57). It predicted that mutating Gly260 and Asp288 to Cys disulfide-linked cyclic site 2 peptide of β3 does not affect the conformation of the original site 2 peptide sequence QPNDGSHVGSDNHYSASTTM in the 3D structure. We found that the cyclic site 2 peptide bound to CX3CL1 and sPLA2-IIA to a similar extent to non-cyclized b3 site 2 peptides in ELISA-type assays (data not shown).

### Binding of site 2 peptide to CCL5

Wells of 96-well Immulon 2 microtiter plates (Dynatech Laboratories, Chantilly, VA) were coated with 100 μl PBS containing CCL5 for 2 h at 37°C. Remaining protein binding sites were blocked by incubating with PBS/0.1% BSA for 30 min at room temperature. After washing with PBS, GST-tagged cyclic site 2 peptide was added to the wells and incubated in PBS for 2 h at room temperature. After unbound GST-tagged site 2 peptides were removed by rinsing the wells with PBS, bound GST-tagged site 2 peptides were measured using HRP-conjugated anti-GST antibody and peroxidase substrates.

### Binding of soluble αIIbβ3 to chemokines

ELISA-type binding assays were performed as described previously (5). Briefly, wells of 96-well Immulon 2 microtiter plates (Dynatech Laboratories, Chantilly, VA) were coated with 100 μl PBS containing chemokines for 2 h at 37°C. Remaining protein binding sites were blocked by incubating with PBS/0.1% BSA for 30 min at room temperature. After washing with PBS, soluble recombinant αIIbβ3 (AgroBio, 1 μg/ml) was added to the wells and incubated in HEPES-Tyrodes buffer (10 mM HEPES, 150 mM NaCl, 12 mM NaHCO_3_, 0.4 mM NaH_2_PO_4_, 2.5 mM KCl, 0.1% glucose, 0.1% BSA) with 1 mM MnCl_2_ for 1 h at room temperature. After unbound αIIbβ3 was removed by rinsing the wells with binding buffer, bound αIIbβ3 was measured using anti-integrin β3 mAb (AV-10) followed by HRP-conjugated goat anti-mouse IgG and peroxidase substrates.

### Activation of soluble αIIbβ3 by chemokines

ELISA-type binding assays were performed as described previously (14). Briefly, wells of 96-well Immulon 2 microtiter plates were coated with 100 μl PBS containing γC390-411 for 2 h at 37°C. Remaining protein binding sites were blocked by incubating with PBS/0.1% BSA for 30 min at room temperature. After washing with PBS, soluble recombinant αIIbβ3 (1 μg/ml) was pre-incubated with chemokines for 10 min at room temperature and was added to the wells and incubated in HEPES-Tyrodes buffer with 1 mM CaCl_2_ for 1 h at room temperature. After unbound αIIbβ3 was removed by rinsing the wells with binding buffer, bound αIIbβ3 was measured using anti-integrin β3 mAb (AV-10) followed by HRP-conjugated goat anti-mouse IgG and peroxidase substrates.

### Activation of cell-surface αIIbβ3 by chemokines

αIIbβ3-CHO cells were cultured in DMEM/10% FCS. The cells were resuspended with HEPES-Tyrodes buffer/0.02% BSA (heat-treated at 80°C for 20 min to remove contaminating cell adhesion molecules). The αIIbβ3-CHO cells were then incubated with chemokines for 30 min on ice and then incubated with FITC-labeled γC390-411 (50 μg/ml) for 30 min at room temperature. The cells were washed with PBS/0.02% BSA and analyzed in BD Accuri flow cytometer (Becton Dickinson, Mountain View, CA) or Attune flow cytometer (ThermoFischer Scientific, Waltham, MA). For time-course studies, FITC-labeled γC390-411 was added to cell suspension first and the mixture was kept on ice, and then incubated with chemokines for the time indicated before analyzed in flow cytometry. The data were analyzed using FlowJo 7.6.5.

### Statistical analysis

Treatment differences were tested using ANOVA and a Tukey multiple comparison test to control the global type I error using Prism 7 (GraphPad Software).

## Acknowledgments

We thank Dr. Barry Collar (Rockefeller University) for insightful discussion, and Dr. Sandy Shattil (University of California San Diego) for reagent. We also thank Dr. Samuel Hwang (UC Davis) for the access to flow cytometry.

## Author contributions

Y. K. T. performed the experiments. Y. T. conceived the project, designed the experiments, analyzed the data, and wrote the article. M.F. analyzed the data, discussed, and edited the article.

## Funding and additional information

This project was supported by Pilot funding from Comprehensive Cancer Center at UC Davis School of Medicine. This work is partly supported by the UC Davis Comprehensive Cancer Center Support Grant (CCSG) awarded by the National Cancer Institute (NCI P30CA093373).

## Conflict of interest

The authors declare that they have no conflicts of interest with the contents of this article.

## Notes

### Competing Interest Statement

The authors have declared no competing interest.

